# Elimination of peripheral auditory pathway activation does not affect motor responses from ultrasound neuromodulation

**DOI:** 10.1101/428342

**Authors:** Morteza Mohammadjavadi, Patrick Peiyong Ye, Anping Xia, Julian Brown, Gerald Popelka, Kim Butts Pauly

## Abstract

Recent studies in a variety of animal models including rodents, monkeys, and humans suggest that transcranial focused ultrasound (tFUS) has considerable promise for non-invasively modulating neural activity with the ability to target deep brain structures. However, concerns have been raised that motor responses evoked by tFUS may be due to indirect activation of the auditory pathway rather than direct activation of motor circuits. In this study, tFUS-induced electromyography (EMG) signals were recorded and analyzed in wild-type (WT) normal hearing mice and two strains of genetically deaf mice to examine the involvement of the peripheral auditory system in tFUS-stimulated motor responses. In addition, auditory brainstem responses (ABRs) were measured to elucidate the effect of the tFUS stimulus envelope on auditory and motor responses. We also varied the tFUS stimulation duration to measure its effect on motor response duration. We show, first, that the sharp edges in a tFUS rectangular envelope stimulus activate the peripheral afferent auditory pathway and, second, that smoothing these edges eliminates the auditory responses without affecting the motor responses in normal hearing WT mice. We further show that by eliminating peripheral auditory activity using two different strains of deaf knockout mice, motor responses are the same as in normal hearing WT mice. Finally, we demonstrate a high correlation between tFUS pulse duration and EMG response duration. These results support the concept that tFUS-evoked motor responses are not a result of stimulation of the peripheral auditory system.

## Introduction

Transcranial focused ultrasound (tFUS) is a promising technique for non-invasive neural modulation, with spatial resolution superior to transcranial magnetic stimulation (TMS), transcranial direct current stimulation (tDCS) and transcranial alternating current stimulation (tACS) (Bystritsky *et al*., 2011, Yoo *et al*., 2011). In addition, tFUS has potential for being able to reach subcortical structures deeper in the brain compared to these other transcranial modes of stimulation.

Harvey (Harvey, 1929) first demonstrated almost 90 years ago that ultrasound can activate nerve and muscle fibers. Fry (Fry *et al*., 1958) documented 60 years ago that visual-evoked potentials could be suppressed by ultrasound neuromodulation in cats. More recently, Tyler *et al*. (Tyler *et al*, 2008) demonstrated that pulsed ultrasound triggers sodium and calcium ion transport by activating mechanosensitive ion channels in mouse hippocampal CA1 neurons. Menz *et al*. (Menz *et al*., 2013) demonstrated reproducible activation of salamander retina to US stimulation in an isolated retina preparation. Modulation of the neural activity from ultrasound stimulation has been demonstrated in C.elegans nematodes (Kubanek *et al*., 2018), and in several mammals including pigs (Dallapiazza et al., 2018), sheep (Lee *et al*., 2016), non-human primates (Deffieux *et al*., 2013, Wattiez *et al*., 2017) and humans (Lee *et al*., 2016). Transcranial ultrasound stimulation in rodents has resulted in motor responses such as whole body twitches (Tufail *et al*., 2010, King *et al*., 2013, Ye *et al*., 2016). There also is evidence that ultrasound can be applied to modulate region-specific brain activity in rabbits (Yoo *et al*., 2011) and to induce lateralized motor responses in mice (Kamimura *et al*., 2016).

While these studies have been very encouraging for the development of tFUS as a useful technique, concerns have been raised by several that ultrasound motor neuromodulation results may have been confounded by activation of the auditory pathway in at least some animals. Indeed, it has long been known that ultrasound can stimulate the peripheral auditory system. Foster and Wiederhold (Foster and Wiederhold, 1978) observed cochlear and auditory nerve responses from US stimulation directly applied to the dura in cats. They postulated that transients in ultrasound radiation pressure can cause skull resonances in the auditory frequency range leading to activation of the cochlea. More recently, Sato *et al*. (Sato *et al*., 2018) used GCaMP modified mice with thinned skulls and observed auditory cortex activation in response to ultrasound stimulation that was targeted to induce motor responses. They postulated that a pulsed ultrasound waveform contains broadband acoustic frequency components within the hearing range of mice and that the observed motor responses are an auditory startle-like reflex rather than direct activation of central motor neural circuits. Guo *et al*. (Guo *et al*., 2018) used a multielectrode array and observed auditory and somatosensory cortical activity from transcranial pulsed ultrasound stimulation in guinea pig. They showed that transection of the auditory nerves or removal of cochlear fluids eliminated the US-induced activity. These important studies raise the possibility of auditory system activation from certain US stimuli, but it is not clear from these results how much audible sound reaches the auditory system during transcranial ultrasound stimulation. It also is not clear what mechanism is involved with motor response activity, whether the source of the activity is peripheral or central, or how the observed US-evoked sensory side-effects can be reduced or eliminated.

Our overall goal in this study was to identify and understand the sources and mechanisms of auditory system activation from tFUS stimulation applied to elicit motor responses. We performed a series of experiments that eliminated cochlear input with genetically deaf mice, reduced broadband spectral components in the tFUS signal by smoothing the rectangular waveform envelope, characterized direct physiologic measures of auditory function with advanced comprehensive auditory brainstem responses (ABR), and quantified motor activity with detailed electromyography (EMG) measures. We used TRIOBP mutant mice with a targeted allele that are profoundly deaf (Kitajiri *et al*., 2010). We also used mutant Samba LOXHD1 mice with a mutation in LOXHD1 that causes them to be profoundly deaf (Grillet *et al*., 2009).

We showed that tFUS stimuli can indeed produce auditory system activity in normal hearing mice based on ABR responses. However, we also demonstrated that the ABR magnitude from an ultrasound pulse is less than the ABR magnitude from an acoustic broadband signal near auditory threshold in the form of a sound click at 40 dB SPL. We further demonstrated that the ultrasound waveform envelope can be modified to both eliminate or reduce the auditory frequency components as well as eliminate the ABRs in WT normal hearing mice, and that even when using these modified waveform envelopes, the motor response persists with no reduction. Moreover, we showed that there is a direct correlation between ultrasound pulse duration and muscle EMG response duration, supporting the concept that tFUS directly stimulates central motor pathways for US-evoked motor responses rather than relying on auditory system activation.

## Methods and Materials

### Animals, Preparation and Anesthesia

Thirty-two mice were studied. Twenty-one were normal hearing (WT C57BL/6) mice (Charles Rivers, Wilmington, MA, USA) and eleven were genetically deaf knockout mice. Of the 11 deaf mice, 7 were mutant homozygous TRIOBP mice with a targeted allele that causes them to be profoundly deaf because the stereocilia on the inner hair cells in the cochlea fail to form rootlets, allowing the hair cells to be more easily deflected and subject to damage (Kitajiri *et al*., 2010). The remaining 4 deaf mice were mutant homozygous Samba LOXHD1 mice with a mutation in LOXHD1 and are profoundly deaf shortly after birth because of mechanosensory deficits in the inner ear hair cells (Grillet *et al*., 2009). All mice were females 8 – 12 weeks old with a mean body weight of 22 g (+/−4g). All animal procedures were approved by the Stanford Administrative Panel on Laboratory Animal Care.

Mice used for ABR recordings were anesthetized by intraperitoneal injection of a ketamine (100 mg/kg) and xylazine (10 mg/kg) cocktail (Xia, Anping., Song, Yohan., Wang, Rosalie., Gao, Simon S., Clifton, Will., Raphael, Patrick., Chao, Sung-il., Pereira, Fred A., Groves, Andrew K., Oghalai, 2013). An additional dose of anesthetics (25% of the initial ketamine-xylazine dose) was administered as needed to maintain anesthesia level during longer experiments.

Mice used for EMG recordings were anesthetized with a lower dosage intraperitoneal injection of a ketamine (67 mg/kg) and xylazine (6.7 mg/kg) cocktail. After the animal was completely anesthetized, ophthalmic ointment was applied to protect the eyes from drying, and a hair clipper was used to shave the area of the head where the US transducer was coupled. All mice were placed on a heating pad and rectal temperature was monitored and maintained during preparation. After preparation, the animal was placed on an experimental platform such that all four limbs and the tail were suspended.

### Ultrasound Waveform Generation

A 500 kHz ultrasound signal was delivered from an ultrasound transducer (V301, Olympus, Waltham, MA, USA) via either of two waveguides (15 mm and 5 mm aperture). The narrower-aperture waveguide was designed so that subdermal electrodes could be placed without direct contact between ultrasound gel and the electrodes. To characterize the transducer with the waveguides, the end of each waveguide was sealed with polyethylene and filled with degassed water. The voltage traces produced by ultrasound pressure waves in a degassed water tank were recorded with an optical hydrophone (Precision Acoustics, Dorchester, UK) in a transverse plane approximately 2 mm from the tip of the waveguide to approximate the pressure at the estimated location of focal stimulation in the *in vivo* experiments. Hydrophone beam plots for the two waveguides are illustrated in supplemental figures (Figure S1). Using the pressures measured by the hydrophone, spatial peak pulse average intensity I_SPPA_ [W*/cm*^2^] was calculated as:

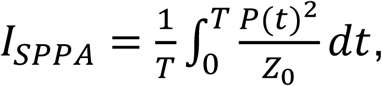

where T = duration [s] of the pressure waveform, P = is pressure [Pa], Z_0_ = is characteristic specific acoustic impedance [Pa s/m] defined as ρc, where ρ = density (1040 kg/m^3^ for brain tissue), and c = speed of sound (1560 m/s in brain tissue) (International Commission on Radiation Units and Measurements 1998).

The spatial-peak temporal-average intensity (I_SPTA_) was defined as

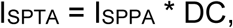

where DC = duty cycle (%) of a pulsed waveform. I_SPTA_ was defined over the duration of the pulsed waveform. Because the mouse skull has negligible attenuation for ultrasound at 500 kHz (Ye *et al*., 2016), attenuation due to the skull was ignored.

Four different ultrasound waveform envelopes were used, each with an overall duration of 80 ms but varying in envelope shape. The spatial peak temporal average intensity, I_SPTA_, was kept constant across all waveform envelopes.

1. Continuous wave with a rectangular envelope (CW_re_). The waveform was generated with a sine wave produced by a function generator (33250 A, Agilent Santa Clara, CA, USA), amplified with a 50-dB radiofrequency amplifier (150 A100 B, Amplifier Research, Bothell, WA, USA) and turned on and off with a rectangular envelope.
2. Pulsed wave with a rectangular envelope and a 1.5 kHz pulse repetition frequency (PRF) (PW_1.5re_). The waveform was generated by triggering the above-referenced function generator with pulse waves produced at the 1.5 kHz pulse repetition frequency using a second function generator (33220 A, Agilent Santa Clara, CA, USA), with an 80% duty cycle rectangular envelope.
3. Pulsed wave with a rectangular envelope and 8.0 kHz pulse repetition frequency (PRF) (PW_8.0re_). This signal is the same as in number 2 above except with a PRF of 8.0 kHz.
4. Continuous wave with a smoothed envelope (CW_se_). The original rectangular envelope was smoothed at both the beginning and the end of the US stimulus with the addition of a radio frequency mixer (ZLW-6, Mini-Circuits, USA). The two coaxial input ports were driven with a sinusoidal continuous wave signal at the center frequency of 500 kHz; and a 90-degree phase shifted single cosine wave. The rapid on and off portion of the rectangular envelope was smoothed over a 12 ms period to reduce the generation of broadband spectral energy (Figure S2).

### Auditory Brain Stem Responses (ABR)

The ABR was measured from 10 mm long platinum 30-gauge subdermal needle electrodes placed 2-3 mm under the skin. The active electrode was placed on the midline of the superior portion of the frontal bone of the skull. The reference electrode was placed at the bottom of the tympanic bulla and the ground electrode was placed on the hindlimb. To establish the hearing sensitivity of both the normal hearing (WT C57BL/6) and one strain (TRIOBP) of the genetically deaf mice, the ABR potentials were measured in the Auditory Core of the department of Otolaryngology at Stanford University as detailed in (Xia, Anping., Song, Yohan., Wang, Rosalie., Gao, Simon S., Clifton, Will., Raphael, Patrick., Chao, Sung-il., Pereira, Fred A., Groves, Andrew K., Oghalai, 2013). Frequency specific acoustic signals were delivered using two high frequency piezoelectric speakers for the ABR measurements (EC1, Tucker-Davis Technologies). The speakers were connected to an earbar inserted into the ear canal and calibrated with a probe-tube microphone (type 4182, Bruel and Kjar, Denmark) inserted through the earbar. The tip of the microphone was within 3 mm of the tympanic membrane. The acoustic ramped sine wave tone pips varied from 4 to 46 kHz for the normal hearing mice and up to 90 kHz for the deaf mice. The sound level at each frequency was raised from 10 to 80 dB SPL(Xia *et al*., 2007). A bioamplifier (DP-311, Warner Instruments, Hamden, CT, USA) was used to amplify the electrode signal 10,000 times and 260 responses were sampled and averaged at each stimulus level following bandpass filtering from 300 to 3000 Hz. The ABR response was considered a valid auditory response if it was 3 times larger than the baseline noise floor.

To establish the hearing sensitivity of the mice used for the motor response experiments, the ABR was measured in a quiet room in the Radiology Sciences Lab of the Department of Radiology at Stanford University. A broadband acoustic click (100 μs rectangular envelope at 40 dB SPL) was delivered in the free field by a loudspeaker positioned 20 cm from the mouse’s right ear. Because there is no difference in ABR between left and right ears (Zheng *et al*., 1999), ABRs were recorded from the right ear only to increase the efficiency of data acquisition. The ABR measurements were acquired with an EEG recording system (SynAmps RT, Compumedics NeuroScan, Australia). The ABR was measured from platinum 10 mm long 30-gauge subdermal needle electrodes. The active electrode was placed 2-3 mm under the skin on the midline of the superior portion of the frontal bone of the skull. The reference electrode was placed below the pinna of the right ear and the ground electrode was placed below the pinna of the left ear. The recorded signals were processed off-line with MATLAB 2017b (Mathworks, USA). The ABRs were derived by averaging 1000 trials following bandpass filtering 300 to 2500 Hz. The ABR response magnitude was calculated as the signal power over a 6 ms period post stimulus onset and determined as present if 5 times larger than the noise floor calculated over an equivalent 6 ms period pre-stimulus onset.

### Electromyography Responses

Motor responses were determined with electromyography (EMG). The EMG recording procedures were similar to previous work (Ye *et al*., 2016). Two 32-gauge enamel-coated copper electrodes were inserted into the triceps muscles of both forelimbs. After attachment of all leads, the US transducer was fixed to a three-axis positioning system. Ultrasound gel was used to couple the waveguide to the head of the mouse.

For all experiments, the transducer coupling cone was positioned approximately 2 mm from the surface of the animal’s head, on the midline 8 mm caudal to the eyes, targeting the midbrain, which contains several motor pathways. A heat lamp distant from the experiment table was used to keep the animal warm. After each mouse experiment, the EMG leads were removed, and the mouse was transferred to an induction chamber for recovery and then returned to its cage.

The EMG signals were amplified with a gain of 1000, and bandpass filtered between 10 Hz and 1 kHz with a preamplifier (World Precision Instruments, Sarasota, FL, USA). Data were acquired at a 2 kHz sampling rate (Lab-Jack U3, LabJack, Lakewood, CO, USA). The sonication parameters were controlled by a computer running software written in MATLAB (Mathworks, Natick, MA, USA).

The EMG signals were post-processed and analyzed using additional software written in MATLAB. For each trial, the DC drift was removed with a 10 Hz Butterworth IIR filter, followed by a notch filter for line frequency (60 Hz) removal. The signal was then full-wave rectified and smoothed with a 15-point moving average filter. A signal was considered representative of a muscle contraction if two conditions were met: first the smoothed EMG signal exceeded a contraction threshold, defined as 6 standard deviations of the signal 100 ms before sonication plus the mean of the signal during this period. Second the smoothed EMG signal exceeded the contraction threshold within a temporal latency less than 200 ms. The contraction latency was defined as the time from the onset of the sonication to the point where the EMG signal exceeded the contraction threshold. The response was then calculated for each set of ultrasound signals as a success rate defined as the number of responses divided by the total number of sonications.

### Experimental design

Four experiments were conducted.

<u>Experiment 1: Motor responses and tFUS level (CW_re_) in hearing and deaf mice</u> The first experiment, designed to compare the EMG motor response to transcranial ultrasound between genetically deaf mice and hearing WT mice, was in two arms. In Experiment 1A, seven hearing WT mice and seven genetically deaf mice (TRIOBP) were used. The ultrasound waveforms used in these experiments were 80 ms (CW_re_) with intensities of 1, 2.79 and 3.78 W/cm^2^. For this experiment, a single run contained twenty sonications in random order: six sonication envelopes at each intensity level, and two sham sonications at 0 W/cm^2^. Ten runs of sonication were applied for each animal, which resulted in 60 sonications at each intensity level. The calculated EMG success rate at each intensity level was averaged over all animals. In the second arm, Experiment 1B, the same experimental setup and ultrasound envelope were used for four hearing WT mice and four deaf Samba (LOXHD1) mice. However, only two levels of intensity 2.79 W/cm^2^ and sham sonications at 0 W/cm^2^ were applied using the narrower-aperture waveguide.
<u>Experiment 2: ABR responses and tFUS envelope in normal hearing WT mice</u> The second experiment was designed to determine whether ABR responses are related to the US envelope and, if so, whether the stimulus waveform could be chosen to minimize or eliminate ABR responses in normal hearing WT mice. Eight mice were sonicated using four US envelopes (CW_re_, PW_1.5re_, PW_8.0re_, and CW_se_) at a constant I_SPTA_ (2.9 W/cm^2^). The ABR was averaged over 1000 sonications for each US envelope.
<u>Experiment 3: Motor responses and US envelope in normal hearing WT and deaf (Samba LOXHD1) mice</u> The third experiment was designed to determine whether motor responses differ with respect to US envelope in normal hearing WT and deaf (Samba LOXHD1) mice. The same experimental setup and US envelopes (CW_re_, PW_1.5re_, PW_8.0re_, and CW_se_, at a constant I_SPTA_ of 2.9 W/cm^2^) were used as those used in Experiment 2. For this experiment, each run consisted of twenty sonications including 4 sham conditions (0 W/cm^2^). Five runs were conducted for each of four animals which resulted in twenty sonications for each US envelope for each animal.
<u>Experiment 4: Motor response and US duration (CW_re_) in hearing mice</u> The fourth experiment was designed to determine if motor response duration was related to sonication duration using a single US envelope (CW_re_) with durations of 80, 160, 320 and 640 ms. The intensity (I_SPTA_ = 2.9 W/cm^2^) was held constant for all US durations. A single run consisted of twenty sonications, four for each of the four durations and four no stimulus sham conditions (0 W/cm^2^) applied in random order. Five runs were conducted in each of six mice.

## Results

### Experiment 1: Genetically deaf mice have similar EMG response to hearing WT mice

We first used ABR measurements to quantify auditory function of both normal hearing WT and both strains of the genetically deaf mice. Figure 1A shows representative ABR waveforms recorded from a normal hearing WT mouse in response to a 16 kHz sound stimulus at decreasing sound pressure levels. Typical peaks at several latencies were observed at high sound levels that decreased in magnitude as sound level decreased. Hearing sensitivity, or threshold, was defined as the lowest acoustic stimulus level at which a response is detectable, in this case, 30 dB SPL at this frequency. Stimuli below the hearing threshold showed no detectable response. The first peak in the ABR waveform also shows an increasing latency as stimulus level decreases, confirming that the responses were from the auditory system. Representative ABR waveforms also are shown for genetic knockout deaf mice, TRIOBP in Figure 1C, and Samba LOXHD1 in Figure 1E. No detectable responses were seen in the deaf mice even at the highest stimulus level, indicating that these mice have no measurable auditory function even for very high-level acoustic stimuli. Mean (and SEM) thresholds as a function of stimulus frequency are shown for hearing WT mice in Figure 1B. These results indicated the typical hearing sensitivity curve for normal hearing WT mice with maximum sensitivity (lowest thresholds) in the frequency range between 8 and 32 kHz and gradually decreasing sensitivity (higher thresholds) for lower and higher frequencies. Similar values as a function of stimulus frequency are shown for the knockout deaf mice, TRIOBP in Figure 1D and Samba LOXHD1 in Figure 1F. In contrast to the results for hearing mice, the deaf mice show no measurable ABR responses in the frequency range between 4 and 90 kHz up to the maximum stimulus level (80 dB SPL).

**Figure 1.**
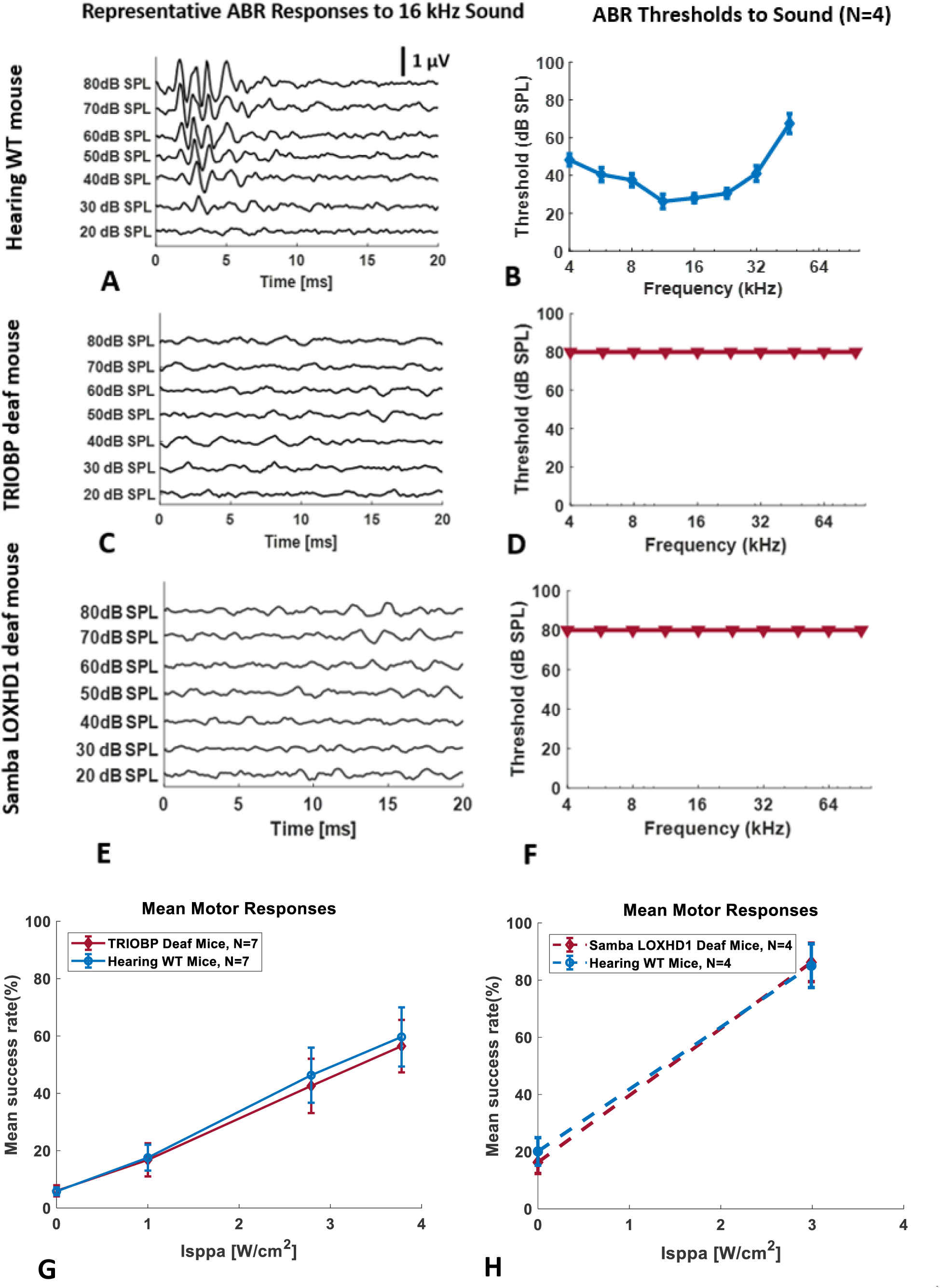
Representative auditory brainstem responses (ABR) to a 16 kHz pure tone at several levels for a normal hearing WT mouse (A), a genetically deaf TRIOBP knockout mouse (C) and a genetically deaf Samba LOXHD1 knockout mouse (E). Hearing threshold is the lowest sound pressure level that produces a detectable response (e.g., 30 dB SPL in A). Mean hearing thresholds (SEM) as a function of frequency for normal hearing WT mice (B), genetically deaf TRIOBP knockout mice (D) and genetically deaf Samba LOXHD1 knockout mice (F). Triangles indicate no response at the highest level used. Mean (SEM) forelimb motor responses (mean EMG success rate in %) as a function of ultrasound (continuous wave rectangular envelope) intensity for normal and TRIOBP genetically deaf mice (G) and for normal hearing and genetically deaf Samba LOXHD1 mice (H).

After confirming that mice with intact hearing have normal auditory function and that both strains of the deaf mice have no measurable auditory function, we then measured motor responses elicited with ultrasound stimuli in normal hearing WT mice and the two strains of knockout deaf mice. Mean (SEM) motor responses (success rate in %) from the EMG in forelimb to a continuous wave US stimulus with a rectangular envelope (CW_re_) as a function of US intensity (Isppa) are shown for normal hearing WT mice and the TRIOBP deaf mice in Figure 1G and for normal hearing WT mice and the LOXHD1 deaf mice in Figure 1H. A two-way analysis of variance (ANOVA) indicated no significant difference in mean motor responses between the normal hearing WT and TRIOBP deaf mice (*p* = 0.718), nor between the normal hearing WT and Samba LOXHD1 deaf mice. Motor responses for a US stimulus that also activates the auditory system in hearing WT mice are identical to motor responses for the same US stimulus in genetically deaf mice.

### Experiment 2: The broadband frequency components of the US stimulus can activate the auditory system

We used ABR measures to determine how US stimulus envelope variations affect auditory activation. Figure 2 shows representative ABR waveforms (voltage as a function of time after US stimulus onset) in a normal hearing WT mouse for four ultrasound (500kHz) signals at a constant pulse-average intensity (I_SPTA_ of 2.9 W/cm^2^) and constant duration (80 msec) but with different envelopes (blue), PW_8.0re_ in Figure 2A, PW_1.5re_ in Figure 2B, CW_re_, in Figure 2C and CW_se_ in Figure 2D. All US signals with a rectangular envelope elicited an ABR at the sharp on portion of the envelope (Figure 2A, B, C). All but one of the US signals with a rectangular envelope elicited an ABR at the sharp off portion of the envelope as well (Figure 2A, C). The sharp off portion of the envelope for PW_1.5re_ did not elicit a detectable ABR (Figure 2B). Larger ABRs were observed for the rectangular envelope pulsed waveforms (PW_1.5re_ and PW_.8.0re_) compared to the rectangular envelope continuous waveform (CW_re_). No ABRs were observed at either the on or off portion of the US signal with a smoothed envelope (CW_se_) or for the no-stimulus sham condition.

**Figure 2.**
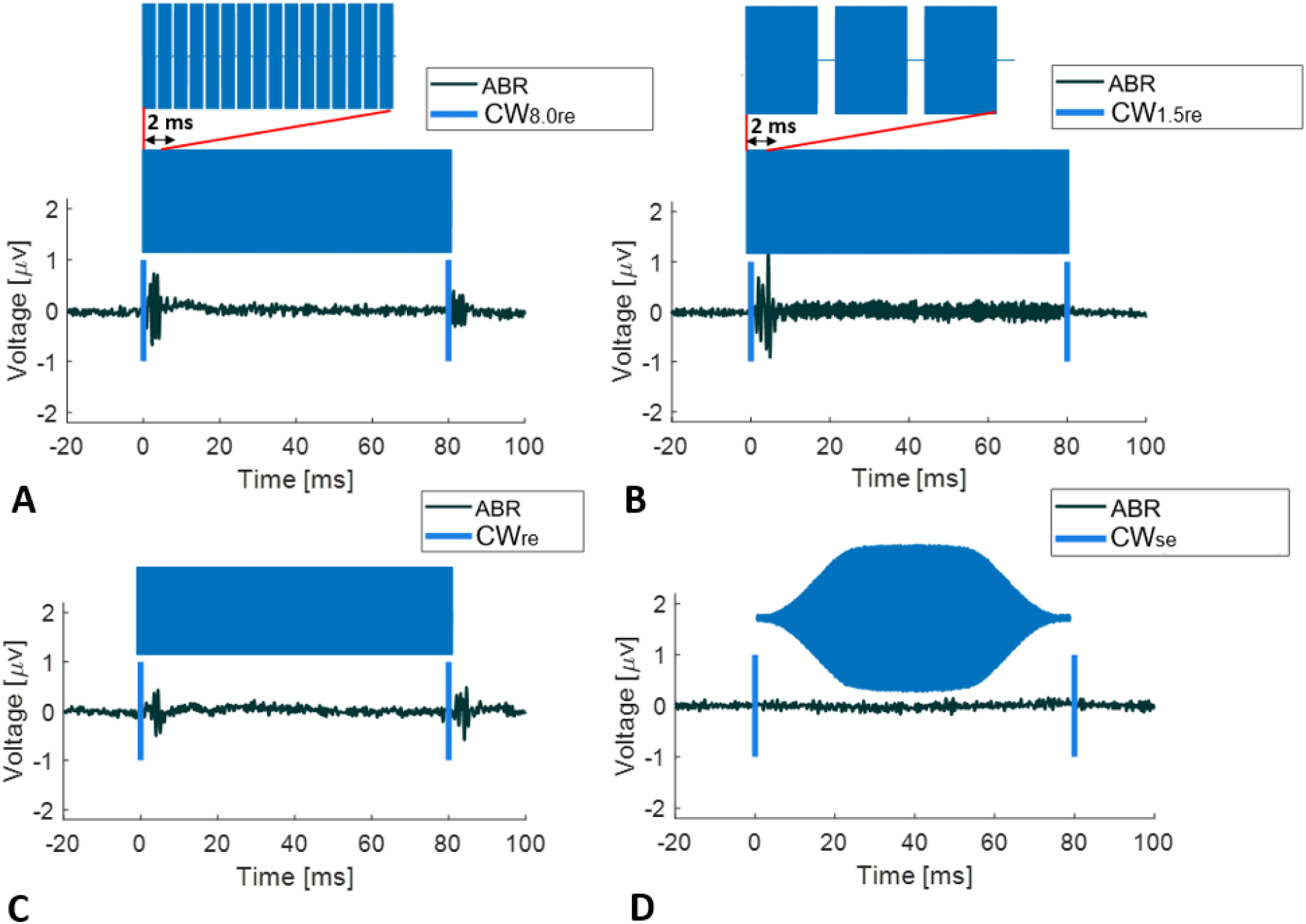
Representative auditory brainstem responses (black wave forms) for an ultrasound signal (500kHz with envelope indicated in blue shading and on and off indicated with blue vertical lines) at equal intensity (2.9 w/cm2) and equal duration (80 msec) in a normal hearing WT mouse. The transcranial ultrasound stimuli were pulsed wave (PW) with rectangular envelope and an 8 kHz pulse repetition frequency (PW8.0re) (A), a pulsed wave rectangular envelope and a 1.5 kHz pulse repetition frequency (PW1.5re) (B), continuous wave (CW) with a rectangular envelope (CWre) (C) and continuous wave with a smoothed envelope (CWse) (D).

We next measured the ABR results at the onset of six stimulus conditions in normal hearing WT mice: an auditory stimulus with a rectangular envelope (100 μs broadband acoustic click at 40 dB SPL, just above threshold), four US stimuli at a constant I_SPTA_ (2.9 W/cm^2^) and a constant duration (80 ms) but with different envelopes (CW_re_, PW_1.5re_, PW_8.0re_, and CW_se_) and a no stimulus sham condition. Figure 3A shows a representative ABR (voltage as a function of time after stimulus onset with stimulus envelope in blue) for each condition. Large ABRs were observed for all signals with a rectangular envelope, the largest for the auditory signal (sound click) and somewhat smaller ABRs for the three US signals (CW_re_, PW_1.5re_, and PW_8.0re_). No ABR was observed for the US signal with a smoothed envelope (CW_se_) nor for the sham condition. Figure 3B shows the mean (and SEM) ABR responses (response defined as the power averaged over 6 ms) in hearing WT mice (\*\**p* < 0.01). The mean ABR response for each of the rectangular envelope signals, the acoustic click and the US CW_re_, PW_1.5re_, and PW_.8.0re_ stimuli, were statistically different from the mean ABR response for the smoothed envelope US stimulus, CWse. (*p* < 0.01, 2-tailed unpaired *t*-test). The mean ABR response for the smoothed envelope US signal, CWse, was indistinguishable from the result for the sham condition (*p* = 0.97).

**Figure 3.**
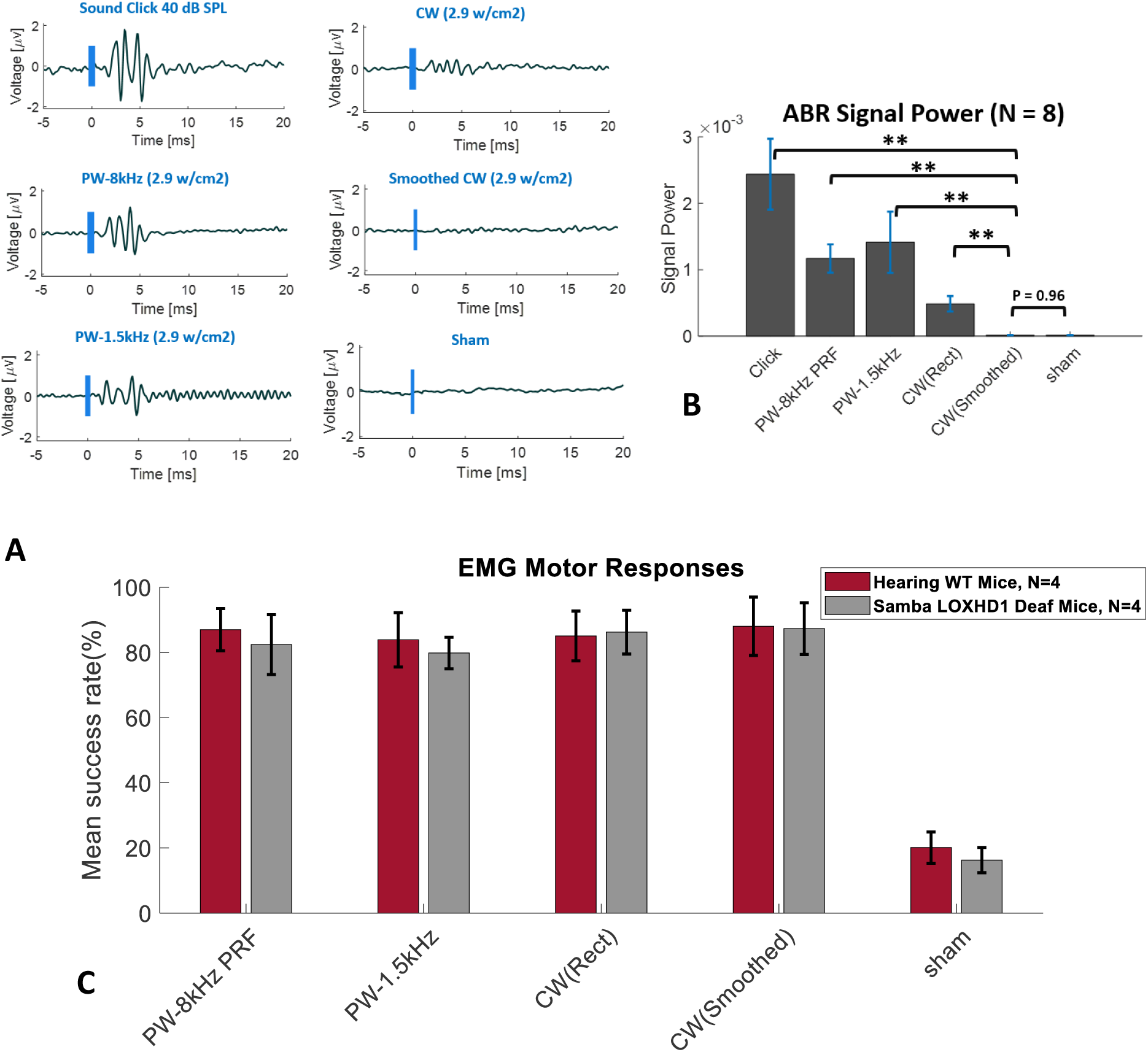
Auditory and motor responses for six stimulus conditions, an auditory stimulus (rectangular envelope broadband acoustic click), four different ultrasound stimuli (CWre, PW1.5re, PW8.0re, and CWse) and a no stimulus sham condition. Representative auditory brainstem response (ABR) waveforms (stimulus in blue) in a normal hearing WT mouse (A). Mean (and SEM) ABR responses (response power averaged over 6 ms) in normal hearing WT mice (** p < 0.01) (B). Mean (SEM) forelimb motor responses (mean EMG success rate in %) for hearing WT (C57BL/6) mice and genetically deaf mice (Samba LOXHD1) (C).

### Experiment 3: Motor responses for different US envelopes are equivalent for normal hearing WT and genetically deaf (Samba LOXHD1) mice

We next investigated motor responses for the US stimuli with different envelopes. Figure 3C shows the mean (and SEM) forelimb EMG motor responses (mean EMG success rate in %) for normal hearing WT mice and for Samba LOXHD1 deaf mice for four US conditions (CW_re_, PW_1.5re_, PW_8.0re_, and CW_se_) at a constant I_SPTA_ (2.9 W/cm^2^) and duration (80 ms) and a no stimulus sham condition. A two-way analysis of variance (ANOVA) indicated no significant difference in mean motor responses between the hearing WT and the Samba LOXHD1 deaf mice for any of the US stimuli (p = 0.635). Motor responses for US stimuli that differ in their ability to activate the auditory system in hearing mice are identical to those in deaf mice.

### Experiment 4: Motor response duration is highly correlated with US duration, but motor response latency is constant and greatly exceeds startle reflex latency

The experiments in Figure 4 were designed to examine how motor responses are affected by US stimulus duration in normal hearing WT mice. A continuous wave rectangular envelope (CW_re_) US stimulus with a constant temporal peak intensity of 2.9 W/cm^2^ was used with durations that varied from 80 ms to 640 ms (80,160, 320 and 640 ms), plus a sham no stimulus condition. Figure 4A shows representative EMG data with stimulus duration indicated in blue. Figure 4B shows the mean (and SEM) forelimb EMG durations as a function of US duration. EMG duration was linearly correlated with US duration (R^2^ = 0.98, *p* < 0.01). Figure 4C shows the mean (and SEM) EMG latencies (the time from the beginning of the US pulse to the beginning of the EMG response) for the four US durations. The mean EMG latencies were constant, around 80 msec, regardless of US duration. Motor response duration is highly correlated with US duration but motor response latency greatly exceeds startle reflex latency (~ 10 msec).

**Figure 4.**
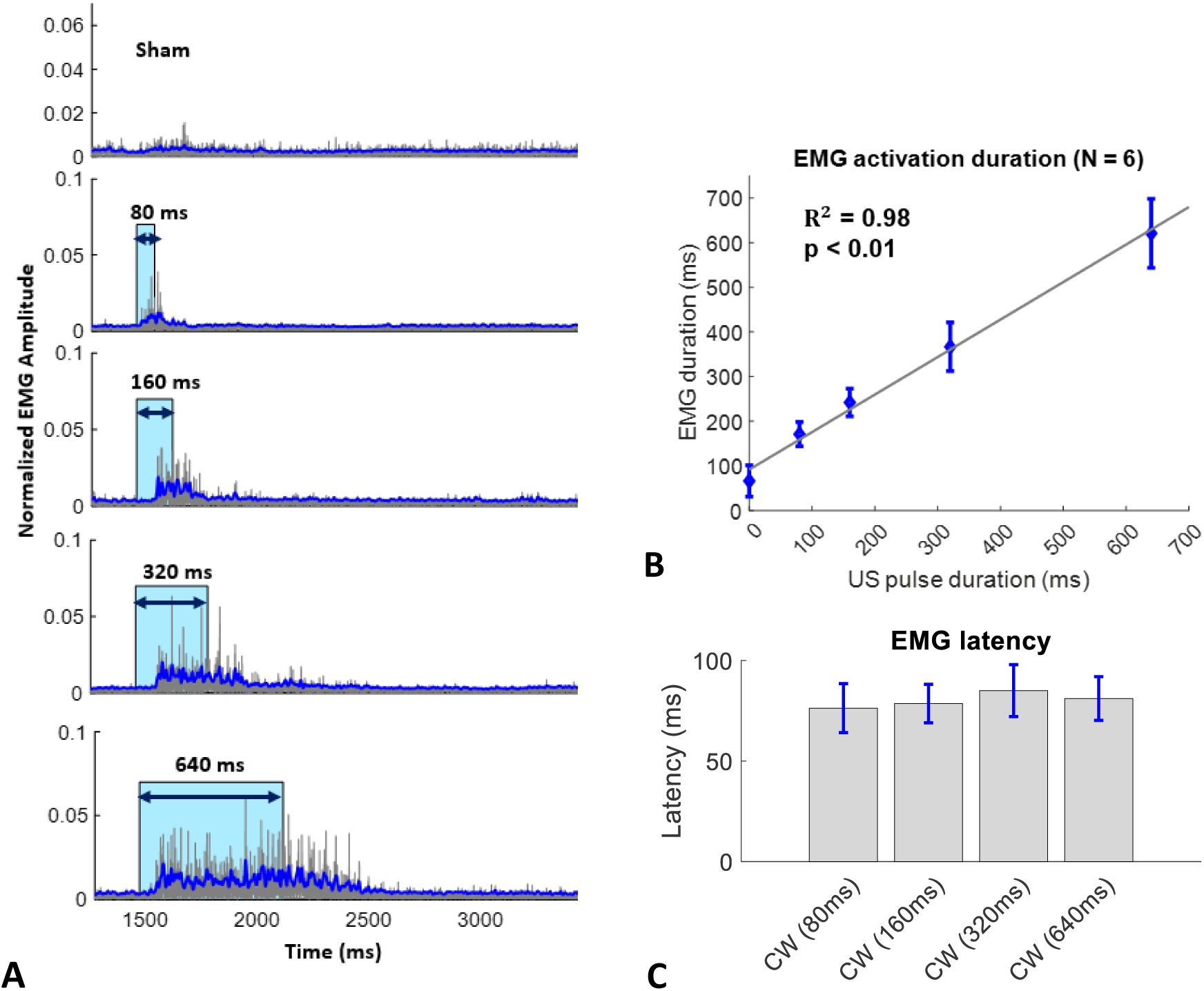
Forelimb muscle activity (EMG) in response to four rectangular envelope continuous wave ultrasound pulse durations and a no stimulus (sham) condition in normal hearing WT mice. Representative EMG waveforms with stimulus indicated in blue (A) in and individual mouse. Mean (and SEM) EMG contraction duration as a function of ultrasound continuous wave (CWre) pulse duration (B). Mean (and SEM) EMG latency for four (CWre) pulse duration (C).

## Discussion

With escalating hopes that ultrasound neuromodulation might become both a new therapeutic tool for the nervous system and a research technique for investigating aspects of brain function, the possibility that motor responses induced by US stimulation could be explained as an auditory artifact has raised some serious concerns. In the absence of a clear understanding of US neurostimulation mechanisms, there remains a critical need to consider alternative explanations that may undermine the possibility of a direct interaction between ultrasound stimulation and nervous system function.

We sought to examine this issue in more detail and determine to what extent US-induced auditory artifacts could be a factor in US neuromodulation experiments. In this study we have corroborated the findings of Foster and Wiederhold, 1978, Sato *et al*., 2018 and Guo *et al*., 2018 by showing that transcranial ultrasound stimulation at a frequency well above the hearing range can be detected by the auditory system in small mammals. Nevertheless, we also have shown that the source of the auditory activation is a portion of the rectangular envelope of the US stimulus waveform that generates vibratory signals in the auditory range that propagate to the sensory cells in the cochlea. We did this, in part, by modifying the ultrasound envelope in such a way as to eliminate most if not all peripheral auditory activation yet while still maintaining the motor response. As further evidence, we took two different strains of genetically deaf mice and confirmed that they exhibited no auditory responses to sound stimuli, yet they maintained the same kind of motor responses as those seen in normal hearing mice. These points, taken together, strongly suggest that the motor response to ultrasound stimuli previously reported in rodents does not rely on activation of the auditory system and is not simply an auditory startle reflex.

A key element of our work is the finding that the ultrasound waveform envelope can contribute to the activation of the peripheral auditory system and thereby produce an ABR response. The broadband frequency components that arise during the near-instantaneous rise and near-instantaneous fall of an US stimulus with a rectangular envelope can indeed activate the cochlea and generate afferent auditory pathway activity as shown by ABR responses. Yet, crucially, we have also shown that smoothing the US waveform envelope by prolonging the rise and fall times can eliminate the ABR response without affecting the expected motor responses (Figure 3B and C).

Interestingly, it is well known that when power is applied at an audible pulse repetition frequency, US transducers can create acoustic signals in the air that are audible to humans. This has raised some questions about a possible acoustic confound or distractor in some US stimulation experiments. Although the authors did not report their reason for doing so in their study (Wattiez *et al*., 2017), Wattiez *et al*. attempted to reduce these air-conducted audible artifacts in their study on non-human primates by using a procedure not unlike ours: they prolonged the rise and fall times at the beginning and the end of each US pulse. However, they chose a much shorter rise and fall time (5 ms) compared with ours (12ms) and they did not report on any comparisons between prolonged rise and fall times and rectangular envelope sonications. We believe our study is the first to demonstrate the clear importance of smoothing the US simulation envelope in eliminating auditory confounds.

Given the benefits of smoothing the US envelope in reducing auditory artifacts, this raises a concern over the use of pulsed US because sequences of pulsed wave rectangular envelopes are likely to exacerbate the problem of generating spectral components in the auditory range compared with a single longer pulse used in continuous waves. In our study we showed that pulsed wave rectangular envelope stimuli do indeed elicit larger ABR responses than continuous wave rectangular envelope stimuli at the same intensity. Because pulsed US has often been preferred over CW US, with some researchers suggesting pulsed US may be more effective than CW (Kubanek *et al*., 2018), further work needs to be performed to determine the theoretical and experimental range of ultrasound stimuli frequency components that result in an acceptably small ABR response to ultrasound yet maintain the desired motor response. A useful study would be to do a more thorough parametric exploration comparing continuous wave and pulsed wave ultrasound stimuli envelopes in animal models to establish a threshold for auditory system activation.

Although our findings agree with Sato *et al* in demonstrating the presence of an audible component when tFUS is applied in certain settings, our results would seem to differ greatly with theirs on the magnitude of the effect. Both studies sought, in part, to compare the effects of tFUS with the effects of audible acoustic stimulation but in different ways, so caution is required in comparing the results. Sato *et al* used a very high level acoustic stimulus (108 dB, presumably SPL) with predominant spectral energy at 1.5 kHz, a frequency range to which rodents are quite insensitive (Heffner *et al*., 2001). In our measurements of hearing thresholds in normal hearing WT mice (Figure 1B), the most sensitive hearing thresholds are in the frequency range between 12 kHz and 20 kHz with worsening thresholds for frequencies both above and below this frequency range. As a result, Sato *et al*. had to increase the amplitude of their 1.5 kHz sound stimuli substantially to render the GCaMP signal similar to that obtained from their broadband tFUS stimuli.

In arguing that ultrasound stimulation may, in part at least, be an auditory startle response, Sato *et al*. also showed that chemical deafening reduced the motor responses to ultrasound. We believe that the loss of motor activity due to chemical deafening could be, at least, partially explained by central neurotoxic effects associated with antibiotic use. In addition to ototoxicity, aminoglycosides have been known to cause peripheral neuropathy, encephalopathy and neuromuscular blockade (Grill and Maganti, 2011, Segal *et al*., 1999, Parsons *et al*., 1992). Aminoglycoside antibiotic-induced neuromuscular paralysis, explained by inhibition of quantal release of acetylcholine in the neuromuscular junction pre-synaptically, has been well documented both clinically and in experimental animals (Parsons *et al*., 1992, Paradelis, *et al*., 1980, Fiekers, 1983). Hence, chemical deafening techniques with aminoglycoisides may not be a reliable control for exploring US-elicited behavioral motor responses.

We believe that future investigations should carefully consider any possible activation of other sensory systems such as vestibular and tactile. It is known that otoconial organs in the peripheral vestibular system are sensitive to low frequency sounds in mice (Jones *et al*., 2010, Yeomans *et al*., 2002). The vestibular system may contribute to the behavioral and cortical responses for frequencies that are below the frequency range of the mouse auditory system, a concept worth investigating.

By eliminating the peripheral auditory system as a source of motor system activation by both smoothing the US waveform and knocking out the cochlear sensory cells, we believe we can argue with greater confidence that ultrasound stimulation can directly modulate central motor neural circuits. Having confirmed that common ultrasound waveforms used for motor system activation can result in peripheral auditory system activation in hearing mice, we postulate that these ultrasound stimuli both directly activate central neural circuits as well as activate the peripheral auditory system. We have shown that by using smoothed US waveforms ultrasound can activate central motor neural circuits in both hearing intact mice and genetically deaf mice with no evidence of peripheral auditory activity.

Our pulse duration results also provide evidence in support of direct activation of central motor neural circuits via ultrasound stimulation rather than via a startle reflex. A startle reflex can be elicited by intense stimulation of the tactile, auditory and/or vestibular systems (Yeomans *et al*., 2002, Li *et al*., 2001) and is defined as a sudden motor movement with a short latency, ≤10 ms (Caeser *et al*., 1989, Cassella *et al*., 1986) in rodents. If the ultrasound stimulus evoked an auditory induced startle response, a motor response would have been observed as a single, short latency, short duration EMG response at the instantaneous rise time, and possibly the instantaneous fall time, of the rectangular envelope of the US stimulus (Figure 2). The EMG motor response latency, ~80 ms, (Figure 4C) was greater than the typical latency of a startle reflex, ≤ 10 ms. Further, the EMG response duration was strongly and linearly correlated with ultrasound pulse duration. These two observations suggest continuous and direct activation of the central motor pathways rather than a startle reflex response.

Finally, adding further weight to the idea that US stimulation acts directly on central motor neural circuits, several other studies have shown some level of stimulus localization with larger transducers and higher ultrasound frequencies, suggesting that smaller focal spots may result in direct neuromodulation effects with superior targeting specificity (Li *et al*., 2016, Kamimura *et al*., 2016, Yoo *et al*., 2011). Further work is needed to determine more precisely which central motor neural circuits are being directly activated by ultrasound, perhaps using implanted EEG or cranial window whole brain measuring techniques.

Although our findings should provide reassurance to researchers investigating the effects of US neuromodulation, studies such as Sato *et al*. and Guo *et al*. are important in questioning the fundamentals of the field. The importance of considering auditory confounds is also relevant to assessing the effectiveness of ultrasound neuromodulation in humans. Reports in recent years have demonstrated the effects of transcranial focused ultrasound on human primary motor cortex activity (Legon *et al*., 2018), primary somatosensory cortex activity (Legon *et al*., 2014, Legon *et al*., 2018) and primary visual cortex activity (Lee *et al*., 2016). However, auditory confounds have been observed in other transcranial methods such as Transcranial Magnetic Stimulation (TMS) where the discharge of the magnetic stimulation coil (Pascual *et al*., 1992) was found to cause permanent shifts in hearing threshold in the unprotected ears of experimental animals (Counter *et al*., 1990). For transcranial ultrasound neuromodulation to become an effective clinical tool, it is therefore important that researchers take into account the auditory phenomenon presented in this and other studies and consider how it can be minimized while still achieving the intended neural modulation.

## Acknowledgment

We would like to thank Anthony Ricci for his helpful discussions and feedback on the auditory results. The authors also thank Nicolas Grillet for donating the genetically deaf Samba LOXHD mice in addition to providing helpful scientific input on the genetic knockout mice. We would also like to acknowledge the Stanford OHNS auditory function core which is supported by the Stanford Initiative to Cure Hearing Loss by a generous gift from the Bill and Susan Oberndorf foundation.

